# Phospholipase D1 and Phosphatidic Acid are required for MVE Fusion and Exosome Secretion

**DOI:** 10.64898/2026.01.21.700874

**Authors:** Melodie T. Nguyen, Broderick L. Bills, Andre C. Allen, Megan L. Hulser, Claire Jiang, Michelle K. Knowles

**Affiliations:** Molecular and Cellular Biophysics Program, University of Denver, Denver, CO 80210; Department of Chemistry and Biochemistry, University of Denver, Denver, CO 80210

**Keywords:** Phospholipase D1, multivesicular endosomes, exosomes, secretion, phosphatidic acid, constitutive exocytosis

## Abstract

Extracellular vesicles (EVs) mediate critical intercellular communication, yet the molecular mechanisms that govern multivesicular endosome (MVE) fusion with the plasma membrane and exosome release remain poorly understood. Phospholipase D1 (PLD1) produces phosphatidic acid (PA), a lipid involved in membrane remodeling, but when and how PLD1 and PA act during exosome secretion has not been defined. Here, we used immunofluorescence and total internal reflection fluorescence microscopy (TIRFM) to track individual CD63^+^ MVEs together with fluorescent PLD1 or a PA reporter (GFP-PASS) in A549 cells. PLD1 localized to CD63^+^ MVEs during visiting, docking, and fusion. Inhibition or knockdown of PLD1 significantly reduced MVE fusion frequency and decreased the number of secreted small EVs, while causing only a modest reduction in vesicle availability at the plasma membrane. PLD1 inhibition also increased the number of membrane-proximal lysosomes, suggesting that MVEs are diverted toward degradation when fusion is impaired. Meanwhile, PA dynamics were stage-specific: PA remained low on visiting vesicles, gradually accumulated during docking, and exhibited a sharp spike followed by loss during fusion. PA has been reported to stabilize negative curvature and potentially play a role during fusion. To test whether PA influences fusion pore behavior, we quantified CD63 decay duration (t_1_/_2_) for individual fusion events. K-means clustering revealed that vesicles with longer decay durations had higher PA levels, whereas short-decay events showed minimal PA. The PA intensity correlated positively with decay duration, while cytosolic GFP did not, indicating a specific relationship between PA and exosome release rates. Furthermore, pharmacological activation of PLD1 increased the proportion of long-decay events. Together, these findings demonstrate that PLD1-generated PA regulates MVE fate at two levels: it promotes the docking-to-fusion transition and prolongs exosome release. This identifies a lipid-based mechanism that controls both the efficiency and kinetics of exosome secretion.

**SIGNIFICANCE STATEMENT:** Exosome release is essential for cell–cell communication in processes such as development, immunity, and cancer. However, the mechanism that determines whether a multivesicular endosome fuses with the plasma membrane to release exosome cargo remains unclear. We identified a lipid-based mechanism that regulates both fusion and the rate of content release. We show that phospholipase D1 (PLD1) and its product phosphatidic acid (PA) act at late stages of exosome secretion: PLD1 promotes the docking-to-fusion transition and PA levels quantitatively correlate with content release rate, suggesting that PA stabilizes the pore. These findings demonstrate that lipids can control secretion frequency and kinetics, providing a new framework for understanding and modulating exosome output.

## INTRODUCTION

Exosomes are small vesicles that circulate in the body, where they carry chemical cargo to communicate from one cell to another. Often this process is upregulated in disease states, where exosomes carry biomolecules that aid in disease progression. Specifically, exosome secretion is involved in cancer metastasis (1–4) and neurodegenerative disorders (5–7) where disease states are thought to spread to the transfer of unhealthy biomolecules to healthy cells (8–12). Exosomes also aid in the function of healthy cells, assisting intercellular communication, maintaining cellular functions such as motility (13–15) and homeostasis (16–19) . Exosomes are released when multivesicular endosomes (MVEs) fuse with the plasma membrane, releasing their intraluminal vesicles (ILVs). Therefore, proteins that modulate the formation of ILVs, the maturation and trafficking of MVEs to the plasma membrane, and the subsequent docking and fusion of MVEs, have the potential to regulate the release of exosomes.

Phospholipase D (PLD) has emerged as a potential regulator of exosome secretion and cancer progression (20–22). The PLD family consists of six isoforms, but PLD1 and PLD2 have been the most extensively studied. Both PLD1 and 2 show a decrease in exosome secretion when inhibited in breast cancer cells (3T1) (21). PLD1 is in puncta that colocalize with acidic compartments, like MVEs and lysosomes (23– 25). Upon inhibition of PLD1, fewer MVEs are observed by electron microscopy, suggesting that one function of PLD1 could be in the formation and maturation process. PLD1 has also been shown to broadly affect neuroendocrine vesicle secretion (26–29) where it is involved with vesicle docking, movement, and fusion typically through the production of PA. However, the mechanism by which PLD1 and its product, phosphatidic acid (PA), control exosome secretion in space and time is not well understood.

PLD converts phosphatidylcholine (PC) to PA, causing the lipid to change its intrinsic shape (from cylindrical and preferring flat membranes to inverse conical and preferring negatively curved surfaces), the overall charge (from neutral to negative), and potential downstream binding partners. Past work has shown that PLD promotes exocytosis through PA production, which stabilizes negatively curved membranes(30–34) such as those observed at a fusion pore or the inward budding that occurs when forming ILVs. PA is produced at docking and fusion sites during dense core vesicle exocytosis(26, 28, 29, 35, 36). Additional regulation occurs through PA interactions; PA binds to the SNARE protein Syntaxin, possibly directly aiding in docking or fusion where Syntaxin has a demonstrated role (37–39). In addition to Syntaxin, PA also interacts with the SNARE-priming ATPase NSF/Sec18, where it modulates priming through direct binding in a yeast vacuole fusion system (40), and with syntenin in MCF-7 cells, where PA affects ALIX-dependent exosome biogenesis(41). There are many places in the trafficking pathway for PLD1 regulation but its role in MVE trafficking is largely unexplored.

In this work, the function of PLD1 and PA in the trafficking, docking and fusing of MVEs and exosome secretion was elucidated using a combination of small extracellular vesicle (sEV) collection, confocal fluorescence microscopy and total internal reflection fluorescence microscopy (TIRFM), where single fusion events were visualized in space and time concurrent with PA and PLD1 imaging. To image single MVEs, CD63 was used as an MVE marker, with a pH-sensitive probe, pHmScarlet. The trafficking, docking and fusion of CD63+ vesicles were tracked concurrent with imaging of GFP-PLD1 or a GFP labelled probe that binds PA (PASS-GFP). Using inhibition and knockdown experiments, our work clearly demonstrates that PLD1 and PA are required for exosome secretion and regulate MVE fusion with the plasma membrane. By noting when and where PLD1 and PA are present during the different stages of exosome secretion, a model for regulation was deduced.

## Methods and Materials

### Cell culture

A549 cells were cultured in Dulbecco’s Modified Eagle Medium (DMEM) (Thermo Fisher Scientific, Waltham, MA, USA) supplemented with 10% fetal bovine serum (FBS; Thermo Fisher Scientific) at 37 °C in a humidified incubator with 5% CO_2_. For imaging experiments, cells were seeded into 4-well glass-bottom plates (Cellvis, Mountain View, CA, USA) and transfected the following day with DNA constructs encoding fluorescently tagged proteins using Lipofectamine 3000 (Thermo Fisher Scientific) according to the manufacturer’s protocol. Plasmid amounts ranged from 25–100 ng/well. The EGFP-PLD1 plasmid was a gift from Jeremy Baskin (42). The GFP–PASS construct was a gift from Guangwei Du (Addgene plasmid #193970) (43). The CD63-pHmScarlet construct was generated by modifying pCMV-Sport6-CD63-pHluorin (Addgene plasmid #130901, (44), replacing the pHluorin tag with pHmScarlet. The pHmScarlet probe was originally developed to track VAMP2-positive vesicle docking and fusion in neuroendocrine cells (45).

### Inhibitor and siRNA treatments

For pharmacological inhibition experiments, cells were treated with 0.013% DMSO as a control, 500 nM PLD1-specific inhibitor VU0155069 (PLD1i; Sigma-Aldrich, St. Louis, MO, USA) or 100 nM pan-PLD inhibitor 5-fluoro-2-indolyl des-chlorohalopemide (PLD1/2i; Sigma-Aldrich) for 30 min at 37 °C prior to imaging.

For knockdown experiments, A549 cells were transfected with Silencer Select siRNA targeting PLD1-siRNA (Assay ID s10637) or PLD2-siRNA (Assay ID 11884), or with a scrambled negative control siRNA (cat. AM4635) (all from Thermo Fisher Scientific), using Lipofectamine 3000 (Thermo Fisher Scientific) according to the manufacturer’s protocol. Cells were maintained for 48–72 h prior to imaging. Knockdown was verified with confocal immunofluorescence imaging.

### Small Extracellular Vesicle (sEV) Isolation

For EV experiments, cells were grown in T75 flasks (CELLTREAT, Life Science Products, Frederick, CO, USA) until reaching 70–80% confluency. (i) *PLD inhibitor experiments:* Cells were treated with the same inhibitor concentrations as above for 24–48 h. (ii) *PLD knockdown experiments:* Cells were transfected with identical siRNA concentrations as above for 48 h. Small EVs were isolated from conditioned media using ExoQuick-TC (Systems Biosciences, Palo Alto, CA, USA) according to the manufacturer’s protocol and quantified using ZetaView Nanoparticle Tracking Analysis (ZetaView PMX-120, Particle Metrix GmbH, Meerbusch, Germany)

### Western Blotting

Following sEV collection, adherent cells were trypsinized (5–10 min) until detached, pelleted (1500 × g, 5 min), washed with TBS, and lysed in RIPA buffer (Thermo Scientific, Cat# 89900) on ice for 30 min. Lysates were homogenized by passing through a syringe, mixed 1:1 with 2× Laemmli buffer containing 5% β-mercaptoethanol, boiled (95 °C, 5 min), and loaded onto 4–15% stain-free gels (Bio-Rad, Cat# 4568083). Proteins were transferred to nitrocellulose membranes (Bio-Rad, Cat# 1620215) using a wet transfer system. Membranes were blocked with 5% BSA in TBS-T (Tris-buffered saline with 0.1% Tween-20) for 2 h at room temperature or overnight at 4 °C, incubated with primary antibodies (Actin [C-2], mouse monoclonal, Santa Cruz, sc-8432) overnight at 4 °C, washed, and incubated with fluorescent secondary antibodies (Santa Cruz, sc-516178) for 4 h at room temperature. Blots were imaged on a ChemiDoc Imaging System (Bio-Rad, Hercules, CA, USA).

### Fluorescence confocal microscopy

For Immunofluorescence assays, A549 cells were fixed in 4% paraformaldehyde (Electron Microscopy Sciences, Hatfield, PA, USA) for 30 min and permeabilized with 0.1% Triton X-100 (Sigma-Aldrich, St. Louis, MO, USA) for 15 min. Cells were blocked with 0.5 mg/mL BSA (Thermo Fisher Scientific, Waltham, MA, USA) for 4 hours at 4 °C, then incubated overnight at 4 °C with mouse anti-PLD1 (Santa Cruz Biotechnology, Santa Cruz, CA, USA) and rabbit anti-CD63 (Abcam, Cat# AB252919-1001) primary antibodies. The following day, cells were incubated for 4 hours at 4 °C with Alexa Fluor 594 conjugated goat anti-mouse and Alexa Fluor 488–conjugated goat anti-rabbit secondary antibodies (Thermo Fisher Scientific, Waltham, MA, USA). For PLD1 knockdown experiments, cells were transfected with siRNA for 48–72 h prior to immunofluorescence staining, as described above.

### Total Internal Reflection Fluorescence Microscopy

A549 cells were imaged in FluoroBrite DMEM (Thermo Fisher Scientific, Waltham, MA, USA) using a TIRF microscope (Nikon Ti-U) equipped with 491 nm and 561 nm lasers, as described previously (46). A 60× 1.49 NA objective, 2.5× magnifying lens, and EMCCD camera (Andor iXon 897, Abingdon, UK) were used in combination with a DualView (Optical Insights, Suwanee, GA, USA) to split red and green fluorescence channels onto the camera via a 565LP dichroic with 525/50 and 605/75 emission filters (Chroma Technology, Bellows Falls, VT, USA). Both color channels were acquired simultaneously, and images were collected in Micro-Manager (47) at 0.108 µm/pixel and 100 ms/frame. Cells were kept at 37 °C on a stage heater during imaging.

### Image Analysis

Image analysis was performed in MATLAB (R2024a, MathWorks, Natick, MA, USA). Time-lapse movies were photobleach-corrected using a custom script to maintain constant overall cell intensity. Visiting, docking, and fusing vesicles were identified using a previously established algorithm (46). Briefly, a difference movie was generated, maximum-projected, bandpass-filtered, and analyzed with a peak-finding algorithm to detect spots exhibiting rapid intensity increases. The locations of these intensity changes in the red channel were used to crop corresponding regions from the photobleach-corrected movies (both green and red channels).

From these cropped movies, intensities were calculated as:

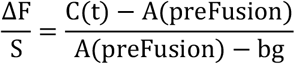

where C(t) is the intensity within a 7-pixel diameter circle in the CD63 channel at the red-channel event location, A is the intensity of a 1-pixel-wide concentric ring separated by a 1-pixel gap from the circle, A(prefusion) is the mean A intensity over the 10 frames preceding fusion onset, and bg is the mean background intensity surrounding the cell.

#### Colocalization

The colocalization of CD63 and PLD1 was measured using a proximity-based approach applied to confocal fluorescence images. First, images of CD63 and PLD1 were processed using a band-pass filter to remove background noise. Vesicles were then detected in each channel using peak-finding algorithms. To determine colocalization, the Euclidean distance between each CD63^+^ vesicle and all PLD1 spots was calculated. A CD63^+^ vesicle was considered colocalized if at least one PLD1 spot was within a defined proximity threshold of 207 nm. The colocalization ratio was computed as the fraction of colocalized CD63^+^ vesicles relative to the total number of CD63^+^ vesicles.

#### K-Means Clustering of Fusion Events by Decay Duration

Fusion event decay durations were calculated using the Full Width at Half Maximum (FWHM) method, where each smoothed intensity trace was used to determine the time interval from peak intensity to the first frame where intensity fell below half-maximum. Events without a valid decay endpoint were excluded, and decay duration longer than 70 frames (7.0 seconds) were also removed to avoid outliers that do not reflect typical fusion behavior. K-means clustering was applied to the resulting decay durations, partitioning the dataset into *k* = 4 groups by minimizing the within-cluster sum of squared distances between data points and their respective cluster centroids. MATLAB’s k-means function was run with 10 replicates and a fixed random seed (rng(1)) to ensure reproducibility, selecting the solution with the lowest total variance. Cluster centroids were sorted in ascending order and labels reassigned for consistent interpretation. This unsupervised approach enabled data-driven classification of fusion events into four kinetic categories, revealing biologically meaningful heterogeneity in post-fusion decay behavior. To compare PMA treated cells, decay durations from all experimental groups (e.g., DMSO, PMA) were concatenated and clustered together using global k-means (k = 4, 10 replicates, Euclidean distance) to ensure consistent cluster definitions across groups. Cluster centroids were sorted in ascending order, with clusters 1–2 designated as short-decay and clusters 3–4 as long-decay events. Group and day identifiers were retained to extract short- and long-decay counts per condition and per day. These counts were compared between conditions using 2×2 contingency tables and Chi-square tests (two-sided), allowing detection of both subtle and pronounced shifts in decay duration distributions.

#### Detection of membrane-proximal vesicles/lysosomes

To identify membrane-proximal vesicles, detection was performed on the average of the first 10 frames from the red fluorescence channel, which enhances the signal of stationary vesicles by reducing frame-to-frame noise. The averaged image was processed using a bandpass filter. Candidate vesicle centers were detected using previously published spot-finding code (48). If a binary cell mask was provided, it was used to restrict detection to within the cell boundary; peaks outside the mask were excluded. The final vesicle positions were visualized by overlaying red circles on the averaged image. To quantify vesicle density, we calculated the number of detected vesicles and divided by the estimated cell area (in μm^2^), using the known pixel size and the total number of pixels within the mask or the image. All work was performed in MATLAB.

## RESULTS

### PLD1 localizes to CD63+ vesicles and is required for exosome secretion

Inspired by the studies demonstrating the essential role of PLD1 in producing PA at stimulated fusion sites (26, 28, 29, 35), we sought to determine if PLD1 is also required for exosome secretion in A549 cells. Confocal immunofluorescence of fixed A549 cells revealed strong PLD1 signal colocalizing with MVEs (Figure 1A), suggesting that PLD1 is in position to potentially regulate exosome secretion. PLD1 was imaged concurrently with CD63, a canonical marker that predominantly labels MVEs (49, 50).To quantify colocalization (Figure 1B), we compared the positions of PLD1 and CD63 puncta. On average, 35% of CD63 spots had a corresponding PLD1 signal (0.352 ± 0.039, mean ± SEM). This colocalization was significantly higher than the negative control, in which the PLD1 primary antibody was omitted (0.004 ± 0.003), but lower than the positive control, where two different anti-CD63 antibodies were used (0.89 ± 0.02). Based on visual observations, it was evident that more PLD1 puncta existed than just those on CD63+ vesicles, suggesting that PLD1 likely marks other types of vesicles too (Figure 1A).

**Figure 1.**
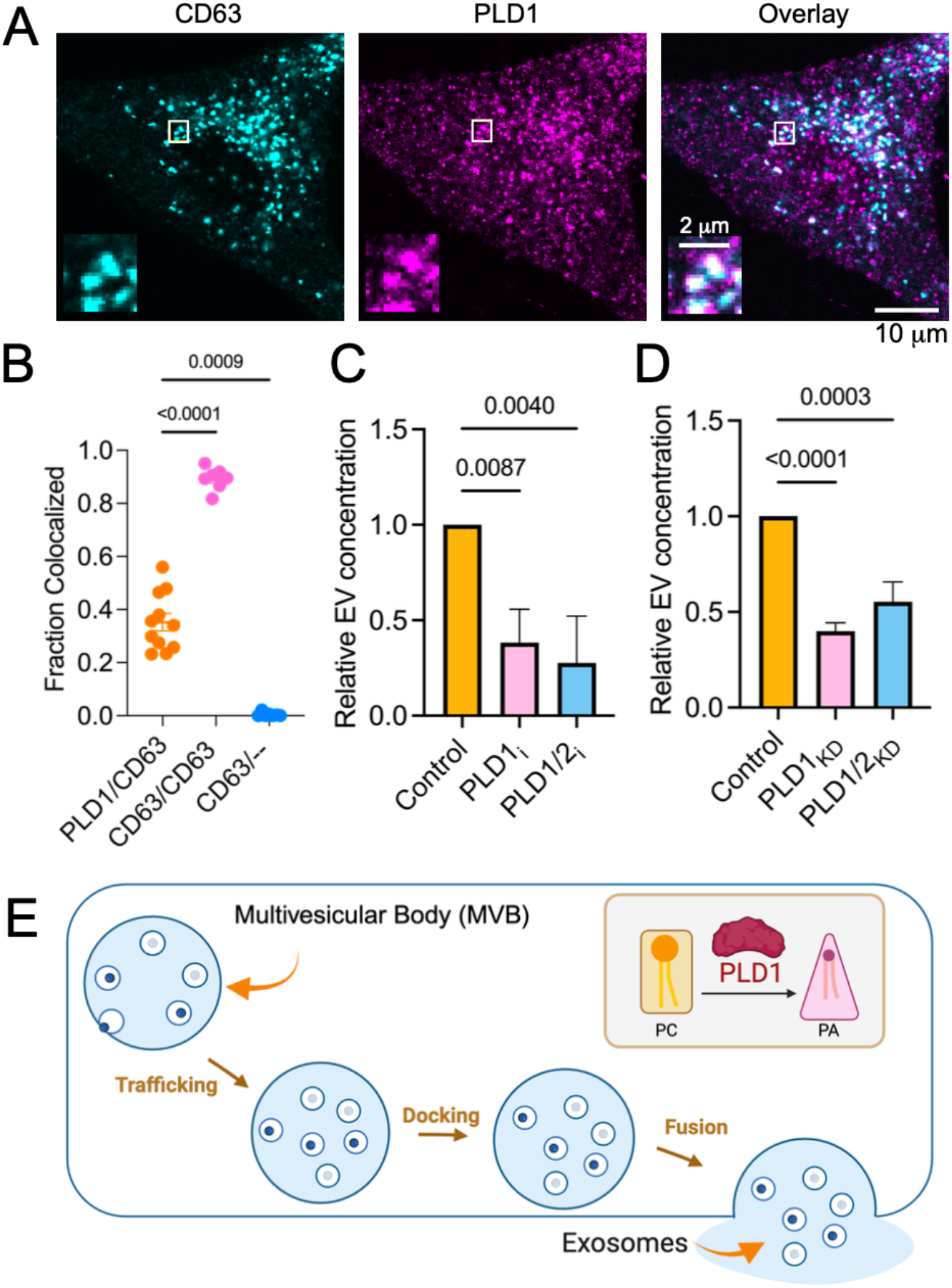
PLD1 is vesicle-localized and required for small EV secretion. **(A)** Confocal microscopy reveals PLD1 colocalization with CD63^+^ vesicles in A549 cells. Representative confocal images of fixed A549 cells stained with anti-CD63 (cyan) and anti-PLD1 (magenta). Overlay image shows extensive colocalization between CD63 and PLD1 signals. Insets highlight vesicles with overlapping fluorescence signals (scale bars:10.0μm and 2.0μm). **(B)** Quantitative colocalization analysis was performed by identifying CD63 and PLD1 fluorescence peaks using MATLAB. The colocalization ratio was defined as the fraction of PLD1 peaks located within 1.5 pixels (207 nm) of a CD63 peak. **(C-D)** PLD1 knockdown or inhibition and small extracellular vesicle (sEV) release. A549 cells were treated with inhibitors or siRNA for 24-48 hours in serum-free media. sEVs (<200 nm) were isolated from conditioned media by precipitation and quantified by nanoparticle tracking analysis (NTA). Vesicle counts were normalized to cell number based on actin levels from Western blotting. Data from three independent experiments were normalized to the control on each day. **(C)** Relative sEV concentration under DMSO (control), PLD1 inhibitor VU00155069 (PLD1i), and dual PLD1/2 inhibition with FIPI. **(D)** Relative sEV concentration under scrambled siRNA (control), PLD1 knockdown (PLD1-KD), and dual PLD1/2 knockdown (PLD1/2-KD). Statistical analysis: one-way ANOVA with Dunnett’s multiple comparisons test. **(E)** Model illustrating key steps in exosome secretion where PLD1 may act; vesicle trafficking, docking, and fusion. (Inset) PLD1 converts phosphatidylcholine (PC) into phosphatidic acid (PA), a lipid involved in membrane remodeling

To assess whether PLD1 function is essential for exosome secretion, PLD1 was either chemically inhibited (Figure 1C) or knocked down by siRNA (Figure 1D), and the concentration of secreted small extracellular vesicles (sEVs) was measured. Inhibiting PLD1 and PLD1/2 using VU0155069 and FIPI, respectively, reduced sEV secretion approximately threefold (Figure 1C), as quantified by nanoparticle tracking analysis (NTA) after collecting sEVs over 48 hours in serum-free media. To account for any effects that the inhibitors had on cell viability, the number of exosomes collected was normalized by the relative Western blot signal intensity of actin (Figure S1A-B). Others have shown a similar reduction with PLD1 and PLD2 specific inhibitors (20, 21, 26). Likewise, siRNA was used to knock down PLD1 and/or PLD2; all siRNA treatments reduced the number of sEVs collected relative to a scrambled RNA control (Figure 1D). Immunostaining confirmed reduced PLD1 of protein expressed after knockdown (Figure S1C-E).

PLD1 inhibition and reduction could alter exosome secretion in a variety of ways, from inhibiting the formation of intraluminal vesicles (ILVs), reducing trafficking to the cell surface, or by reducing fusion.

Figure 1E summarizes each stage where PLD1 could affect the overall number of exosomes secreted. All three are established as potential pathways by which PLD could regulate secretion (20, 36, 51–54) and they are not mutually exclusive. In following sections, we investigate how PLD1 effects each stage.

### PLD1 is present on moving MVEs, increases while MVEs remain docked and is lost post-fusion

To determine which stage of exosome secretion affected by PLD1, we turned to time-resolved, TIRFM methods. Using TIRFM and pH-sensitive fluorescent proteins, as described previously single MVEs were observed moving, docking, and fusing concurrent with GFP-PLD1. To better observe MVEs prior to fusion, CD63-pHmScarlet was designed from the CD63-pHluorin construct, where the pH sensitive fluorescent protein is placed on an extracellular loop of CD63. This places the probe within the MVE and on the surface of the exosomes once secreted. The pHmScarlet probe is slightly visible prior to fusion and clearly shows fusion in cells transfected with both CD63-pHmScarlet and CD63-pHluorin (Figure S3). Using the pHmScarlet probe, MVEs were observed moving, docking and fusing with the plasma membrane. To examine PLD1 dynamics during exosome secretion, we tracked GFP-PLD1 at CD63-positive MVE sites. The relative PLD1 signal was quantified by measuring intensity at the center of the CD63 spot (diameter = 7 pixels), normalized to a surrounding annular background (radius = 11 pixels), as described in the Methods section. The relative signal intensity (ΔF = center – annulus) is then normalized by the cellular expression level (S = annulus – background outside of the cell). Therefore, the ΔF/S is a local intensity or enrichment of the protein at the specific location of the MVE. Time was aligned so that 0 s (“onset”) marks the frame immediately before CD63 intensity begins to rise.

To categorize vesicles into different stages of the secretion process, the CD63 intensity trace was assessed after using an automated MATLAB program to locate locations where a change in intensity is observed (46). Moving vesicles gradually increase and decrease (Figure 2A, purple), whereas docking vesicles gradually increase but remain stationary and do not decrease substantially over time (Figure 2B, dark green), nor does fluorescence increase in the annulus. It is worth noting that a “kiss and run” event, where a pore opens then closes, would have a similar intensity profile. Fusing MVEs increase rapidly (over 200-400 ms), do not move, and the fluorescence radially expands in time as fluorescent CD63 molecules diffuse from the fusion site (Figure 2C, navy blue). After categorizing the MVEs, the PLD1 channel is averaged, and can be assessed for moving, docking and fusing vesicles. Moving MVEs carried abundant PLD1 (Figure 2A, lavender), while docking and fusing vesicles showed lower PLD1 levels (Figures 2B–C, light green and light blue). Cytosolic GFP was used as a negative control and fusion events are shown in Figure 2D; docking and moving events are shown in Figure S4.

**Figure 2.**
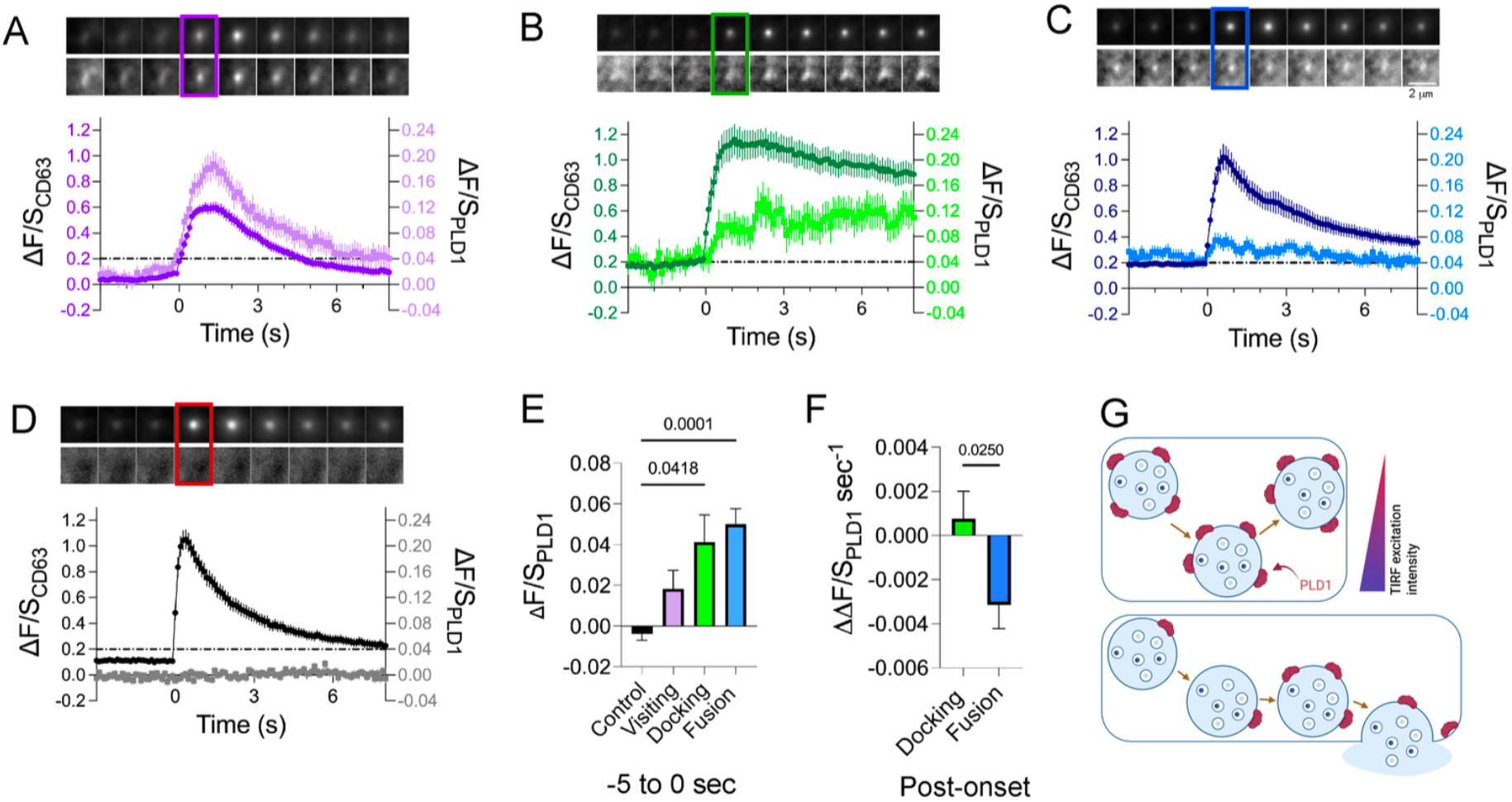
Distinct PLD1 dynamics define vesicle behavior at the plasma membrane. **(A– D)** Representative traces of CD63 (darker shade) and PLD1 or cytosolic GFP (lighter shade) are shown for visiting (A, purple), docking (B, green), fusion (C, blue), and negative control (D, black) vesicles. CD63 (top row) and PLD1 or GFP (bottom row) image montages span from 49 frames before onset to 80 frames after. Each montage frame represents a 15-frame average (1.5 seconds). The boxed regions indicate a focused window spanning 4 frames before to 10 frames after the onset frame (frame 0), highlighting local intensity dynamics during the critical transition period. The full montage captures the broader temporal evolution of CD63 and PLD1 or GFP signals for each vesicle type. Examples were selected from a dataset of 161 visiting, 33 docking, 121 fusion, and 83 negative control events across three independent experiments. **(E)** Mean PLD1 intensity (ΔF/S) during the prefusion phase (49 frames before onset) was significantly higher in docking (n = 33, p = 0.0418) and fusion vesicles (n = 121, p = 0.0001) compared to negative control (n = 83, cytosolic GFP). Visiting vesicles (n = 161) did not differ significantly from control. **(F)** Post-peak PLD1 slopes were significantly different between fusion and docking events (p = 0.0250, Mann– Whitney U test). Fusion vesicles showed a negative slope; docking vesicles displayed a positive slope. **(G)** Schematic model summarizing PLD1 dynamics in each vesicle type. Visiting vesicles display transient and symmetric PLD1 localization mirroring CD63 behavior. Docking vesicles show gradual PLD1 buildup without a sharp peak, while fusion vesicles exhibit a rapid decline in PLD1 following the onset of fusion.

Prior to the onset, moving vesicles contained low PLD1 present, but this increases prior to fusion suggesting that PLD1 accumulates after docking (Figure 2E). The increase in PLD1 over time for MVEs that remain docked was confirmed by evaluating the change in PLD1 intensity after a vesicle docks (Figure 2E, green); the slope (over 10 seconds) increases gradually, whereas the amount of PLD1 decreases post fusion (Figure 2F, blue). Overall, this suggests that PLD1 is present for fusion and on moving vesicles but is displaced post fusion (Figure 2G).

### PLD activity promotes MVE docking and fusion

The accumulation of PLD1 prior to fusion suggests that PLD1 plays a role in the fusion step. To directly determine whether PLD1 activity is essential for fusion, the frequency of MVE fusion events was measured with inhibitors present (Figure 3A) or with the knockdown of PLD1 or PLD1/2 combined (Figure 3A). In both cases, the rate of observed fusion events was significantly reduced, suggesting that the activity of PLD1 and the formation of PA are needed for the fusion process. It is likely that the decrease in fusion events is partly responsible for the decrease in sEVs secreted (Figure 1C-D). Fewer fusion events when PLD1 is inhibited or absent could be due to fewer CD63+ vesicles trafficking to the plasma membrane and/or docked vesicles incapable of fusion. To test if PLD1 is required for trafficking to the plasma membrane, the number of MVEs near the cell surface was evaluated by imaging the surface of live cells using TIRFM. To ensure all CD63+ vesicles were visualized, we neutralized acidic lumens with ammonium chloride to unquench pH-mScarlet, then PLD1 and PLD1/2 were inhibited and compared to a vehicle control. To quantify the imaging data, the number of MVEs in close proximity to the cell surface was measured and normalized by the surface area to evaluate the density of plasma membrane proximal vesicles/μm^2^. We found that membrane-proximal vesicle density was reduced by approximately 20% under PLD1i (*p* = 0.014) and PLD1/2i (*p* = 0.0299) compared to a control (Figure 3B), indicating impaired vesicle accumulation at the plasma membrane when PLD activity is suppressed. Although significant, the reduction is small relative to the large reduction in the frequency of fusion events, suggesting that PLD1 matters more for fusion than for docking.

**Figure 3.**
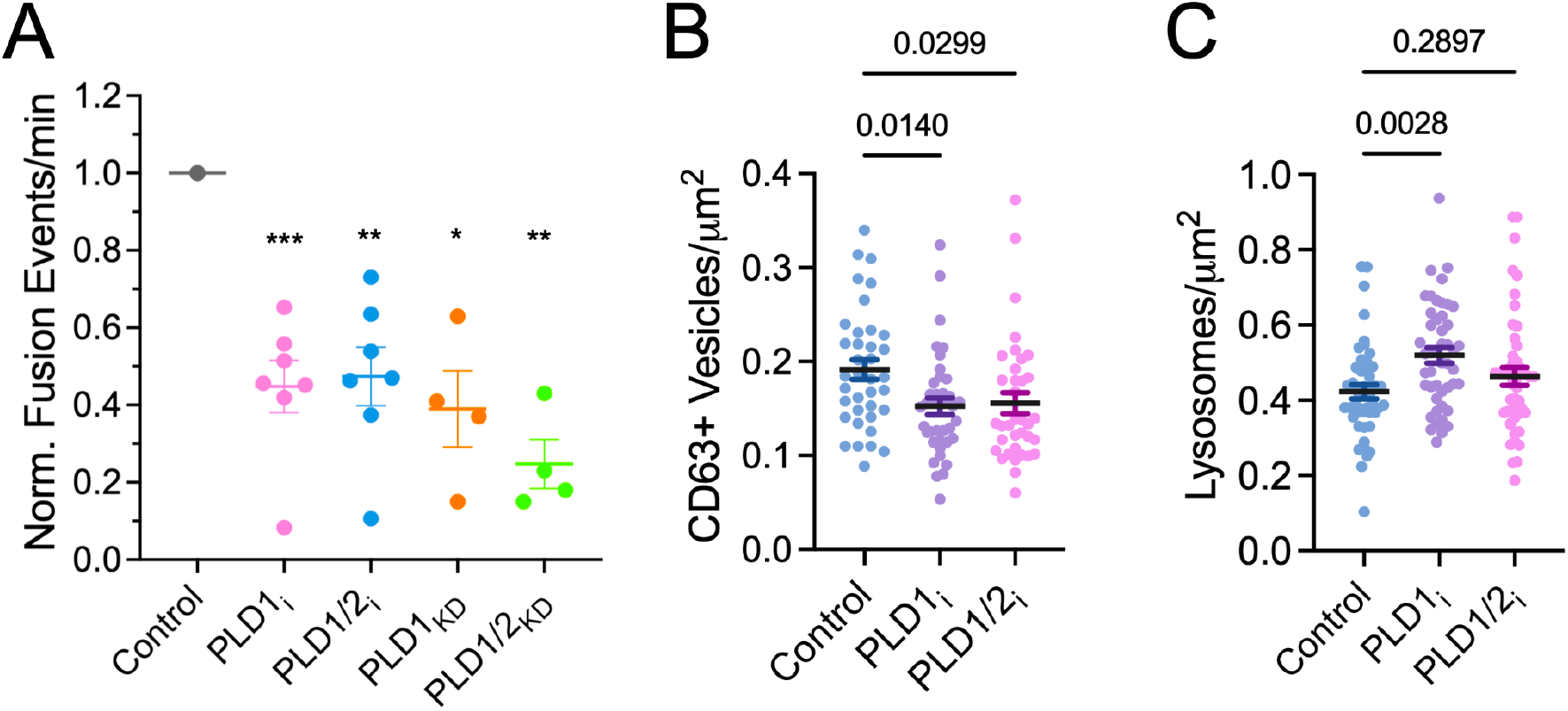
PLD activity regulates fusion frequency and membrane-proximal CD63 vesicle density. **(A)** Normalized fusion event frequencies under pharmacological and genetic modulation of PLD activity. A549 cells were transiently transfected with CD63-pHmScarlet and GFP-PASS, then treated with DMSO (control), the PLD1 inhibitor VU00155069 (PLD1i), or the dual PLD1/PLD2 inhibitor FIPI. Separately, cells were transfected with control siRNA, PLD1 siRNA, or a combination of PLD1 and PLD2 siRNAs. Fusion frequency was calculated as the number of fusion events per 100-second movie, then normalized to the control condition within each experimental day. Fusion frequencies were significantly reduced under PLD1i (\*\*\**p* < 0.001), FIPI (\*\**p* < 0.01), PLD1 knockdown (\**p* < 0.05), and PLD1/2 knockdown (\*\**p* < 0.01) compared to their respective controls. **(B)** Quantification of membrane-proximal CD63 vesicle density at the plasma membrane. A549 cells co-expressing CD63-pHmScarlet and GFP-PASS were treated with 50 mM NH_4_Cl and imaged using TIRFM to enhance visualization. Membrane-proximal vesicles were defined as CD63-positive puncta visible in the TIRF field and detected using a custom MATLAB script employing bandpass filtering and intensity thresholding. Vesicle density was significantly reduced in PLD1i (*p* = 0.014) and PLD1/2i (*p* = 0.0299) conditions compared to DMSO. Each data point represents the vesicle density per cell area across three independent experiments. Cell counts: DMSO, *n* = 37; PLD1i, *n*= 39; PLD1/2i, *n* = 36. **(C)** Quantification of membrane-proximal lysosome density. A549 cells expressing CD63-pHluorin were stained with Magic Red and imaged by TIRFM. Cells were selected based on CD63-pHluorin expression. Membrane-proximal lysosomes were defined as Magic Red-positive puncta and detected using the same MATLAB-based analysis as in (B). Lysosome density was significantly increased under PLD1i (*p* = 0.0028) but not significantly altered under PLD1/2i (*p* = 0.2897) compared to DMSO. Each data point reflects the mean lysosome density per cell from three independent experiments (*n* = 47 for all conditions). Statistical comparisons were performed using *one-way ANOVA*.

### PLD Inhibition Increases Density of Lysosomes near the Cell Surface

MVEs can either fuse with the plasma membrane to release exosomes or merge with lysosomes for degradation (55–59). There is interplay between these fates such that blocking lysosomal degradation increases exosome secretion (55, 60, 61). Notably, phospholipase D enzymes have also been implicated in regulating late endosome fate (41, 62, 63). Therefore, we hypothesized that a reduction in MVE fusion with the plasma membrane would increase the number of lysosomes present. Using the lysosome-specific fluorescent substrate Magic Red and TIRFM imaging, we quantified surface-proximal lysosomes to identify how PLD1 inhibition affects the lysosome density. The number of lysosomes were counted and normalized by the cell surface area. Lysosome density was significantly increased under PLD1i (p = 0.0028), but unchanged with PLD1/2i (p = 0.2897) (Figure 3C). These findings suggest that inhibiting PLD1 shifts vesicle fate away from plasma membrane docking/fusion toward lysosomal degradation.

### Phosphatidic Acid (PA) Localizes to Sites of MVE Docking and Fusion

To investigate the role of PA—the product of PLD1—we used the GFP-tagged PA biosensor PASS (43) to monitor production during MVE activity. Similar to the PLD1 experiments (Figure 2), CD63-pHmScarlet was co-transfected, and vesicles were classified as visiting, docking, or fusion based on their CD63-pHmScarlet profiles (Figure 4A–C). The intensity (ΔF/S) of PASS was measured concurrent with CD63 to determine if PASS is present before, during, or after onset (Figure 4A-C traces). Average montages are shown in the top panels for CD63 and the bottom panels for GFP-PASS. Difference images (“Δ5s”) were generated by subtracting the initial frame (−5 s to 0 s) from the post-onset frame (0 s to 5 s) (Figure 4A–C). In these difference images, a distinct spot is evident for fusion events, but less so for moving and docking vesicles. Prior to onset, docking vesicles appear to target regions of the plasma membrane with detectable PASS signal (Figure 4D), more so than visiting vesicles. Following onset, PASS intensity increases gradually after docking (Figure 4B, light green), suggesting a slow, time-dependent accumulation of PA.

**Figure 4.**
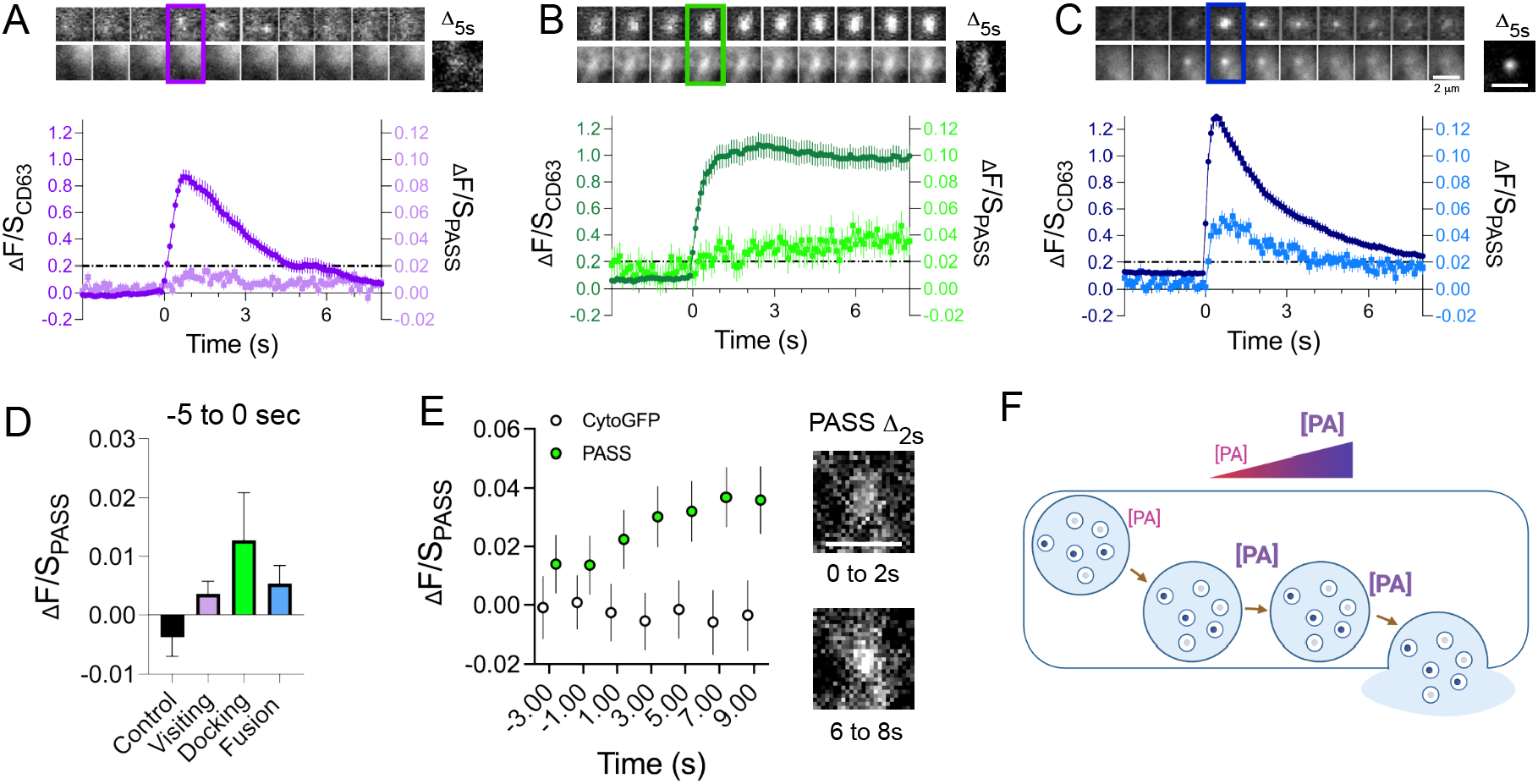
Phosphatidic Acid (PA) accumulation and loss during MVE visiting, docking and fusion. **(A–C)** Representative montage images and intensity traces of CD63 (darker shade) and PASS (lighter shade) are shown for Visiting (A, purple, *n* = 112), Docking (B, green, *n* = 47), and Fusion (C, blue, *n* = 120) vesicles. **Top rows:** CD63 (top) and PASS (middle) montage images span from 49 frames before to 100 frames after onset (frame 0). Each image represents a 15-frame average (1.5 seconds). Boxed regions highlight a focused window from −4 to +10 frames around onset, emphasizing local intensity changes during the transition. **Bottom row:** Full-resolution temporal traces show CD63 and PASS intensity over time for the same vesicle groups, aligned to onset. Difference images (“Δ5s”) were generated by subtracting the average image from −5 to 0 seconds from the image spanning 0 to 5 seconds. **(D)** Quantification of prefusion PASS intensity (−5 to 0 seconds) across vesicle types. Both Docking (*n* = 47) and Fusion (*n* = 120) vesicles exhibited significantly higher PASS intensity than the cytosolic-GFP control (*n* = 83; *p* < 0.0001), whereas Visiting vesicles (*n* = 112) were not significantly different from control (*p* = 0.157). Bars represent mean ± SEM. **(E)** Analysis of PASS dynamics in Docking vesicles. **Left:** Time-binned PASS intensity (green) and cytoGFP control (white) plotted in 2-second intervals. PASS intensity increases significantly over time (linear regression, *p* = 0.0251), while cytoGFP remains flat (*p* = 0.709). **Right:** PASS difference images (Δ2s) showing signal change between 0–2 s (top) and 6–8 s (bottom), to visualize PASS accumulation at docking sites post-onset. **(F)** Schematic model summarizing PA dynamics in each vesicle type. Visiting: PASS signal remains low throughout. Docking: Gradual PASS buildup without sharp peaks. Fusion: Rapid PASS spike following onset, followed by a decline with CD63 loss.

A longer look at docked vesicles (Figure 4E) shows that PASS intensity increases slightly but significantly over 10 seconds. These results indicate that PA is either produced or recruited to MVEs and the nearby plasma membrane. Fusing vesicles, meanwhile, show an increase in PASS immediately post fusion, suggesting that PASS presents on or within MVEs. This signal change may result from vesicle movement into the TIRF field or unquenching of GFP.

### The Amount of PASS is Correlated to the Rate of Exosome Release from Fusion Events

One leading hypothesis as for the role of PA at membrane fusion relates to the inverse conical shape of the lipid itself. With a small headgroup and two fatty acid tails, PA has been shown to support negatively curved membranes (32, 33, 64, 65), suggesting a role at the fusion pore. Therefore, we hypothesized that more PA could stabilize the pore structure and, in doing so, reduce the rate at which content is released from MVEs following fusion.

To assess this, we measured CD63 decay duration for individual fusion events using a full-width-at-half-maximum (FWHM) approach (Figure 5A). For each event, the peak CD63 intensity was identified, and the half-maximum value was calculated. The decay duration was defined as the time from the peak frame to the first frame where CD63 intensity fell to ≤ 50% of the maximum. We then applied k-means clustering—an unsupervised machine learning algorithm that groups data into *k* categories by minimizing within-cluster variance—using only CD63 decay duration as the input feature. This ensured that clustering was determined solely by decay kinetics, without incorporating PA measurements. The algorithm divided the dataset into four clusters (Cluster 1: *n* = 252; Cluster 2: *n* = 141; Cluster 3: *n* = 57; Cluster 4: *n* = 19), ranging from the fastest (Cluster 1) to the slowest (Cluster 4) decay events (Figure 5B). Note that the fusion traces have been aligned to the maximum intensity to be 0s and normalized, different from other figures, to better compare the four clusters, and Figure S4A shows the original fusion traces.

**Figure 5.**
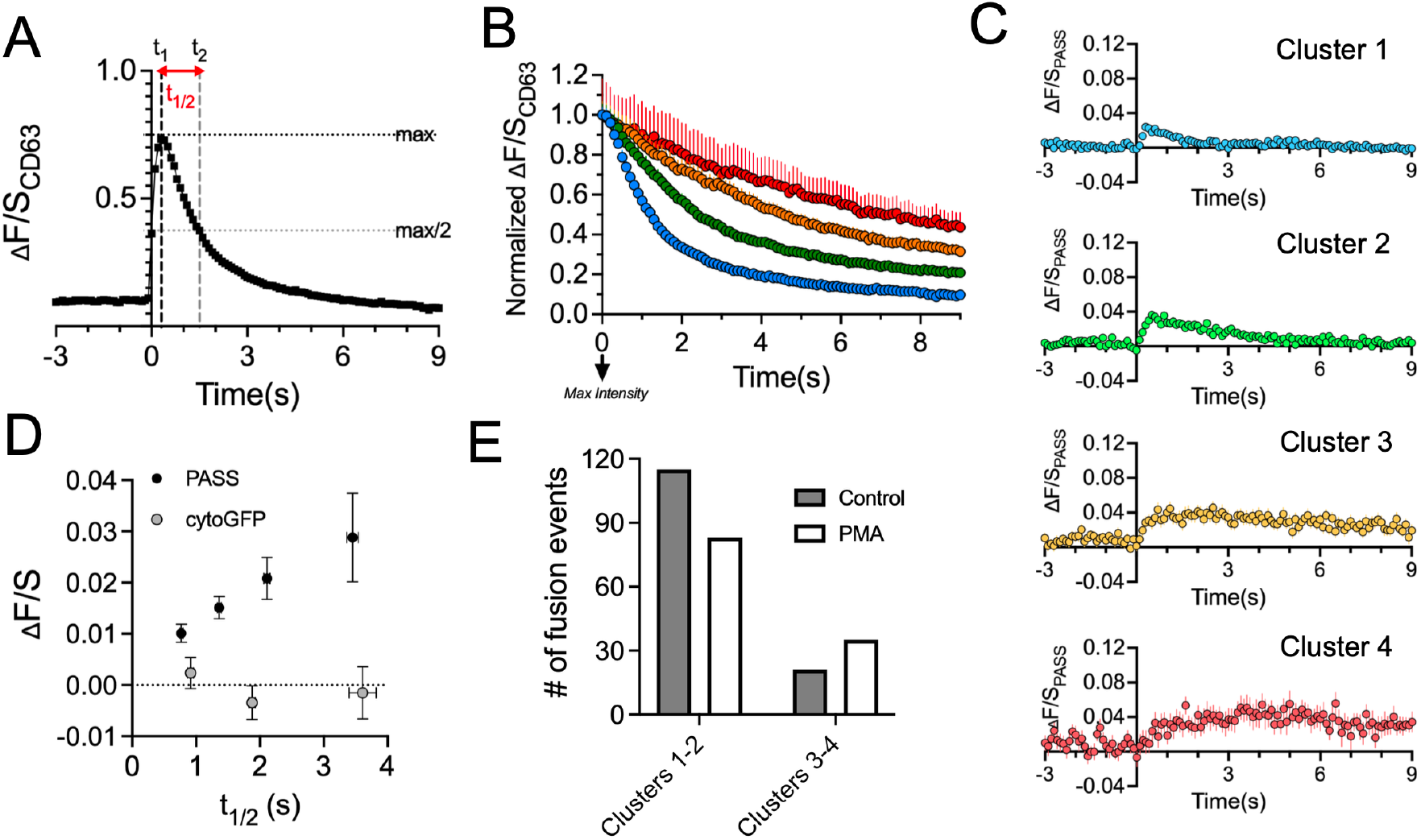
Higher PA levels are associated with prolonged CD63 decay during MVB–PM fusion. **(A)** Diagram illustrating the approach for quantifying CD63 decay duration (t_1/2_). The peak intensity frame (maximum CD63 signal) was identified, and the half-maximum intensity value was calculated. The decay duration was defined as the time interval from the peak frame to the first frame where the CD63 intensity fell to ≤ half of the maximum value. This measurement was performed for each fusion event prior to clustering and used to perform k-means clustering. **(B)** Mean CD63 intensity traces for four k-means–defined clusters (Cluster 1: *n* = 252, Cluster 2: *n* = 141, Cluster 3: *n* = 57, Cluster 4: *n* = 19), aligned such that t = 0 corresponds to the frame of maximum CD63 intensity. Each trace represents the average ± SEM. (**C**) Mean PASS (PA sensor) intensity traces for the same four clusters. Clusters with longer CD63 decay durations tended to maintain higher PA levels throughout the decay phase, whereas short-decay clusters showed lower or rapidly declining PA signal. **(D)** Scatter plot showing the relationship between mean PASS intensity or mean cytosolic GFP intensity and CD63 decay duration for the four clusters. A positive correlation was observed between PASS intensity and decay duration, whereas cytosolic GFP showed no consistent relationship, with an initial negative slope followed by a near-flat trend. **(E)** PMA treatment and activation of PLD1 affects fusion dynamics. The number of events belonging to short-decay clusters (Clusters 1 and 2) and long-decay clusters (Clusters 3 and 4) is shown for DMSO and PMA conditions. A contingency test comparing the distribution of events between DMSO and PMA yielded a statistically significant difference (p = 0.0095), suggesting PLD1 and PA production support longer duration fusion events.

After clustering, we examined the corresponding PASS intensity traces for each cluster (Figure 5C). Clusters 1 and 2 exhibited shorter decay durations and lower PA levels, whereas Clusters 3 and 4 showed prolonged decay and elevated PA throughout the decay phase. Across clusters, mean PASS intensity correlated positively with decay duration (Figure 5D), supporting the conclusion that elevated PA levels are associated with extended decay kinetics. In contrast, cytosolic GFP intensity showed no consistent relationship with decay duration, displaying an initial negative slope followed by a near-flat trend (Figure 5D).

To verify these results and test if more PA yield slower kinetics, we asked if activating PLD1 could increase the number of slow decays (Clusters 3-4). A549 cells were treated with PMA a PKC activator shown to increase PA in cells (66–68). We compared control (DMSO) with PLD activation (PMA) conditions using an independent dataset from that shown in panels A–D. In Figure 5E, the numbers of events in short-decay clusters (Clusters 1 and 2) and long-decay clusters (Clusters 3 and 4) are plotted for DMSO and PMA conditions. PMA treatment shifted the distribution toward long-decay clusters and increased both the per-day fraction of long-decay events and the total number of such events (DMSO: *n* = 21; PMA: *n* = 35; p = 0.0095, contingency test). Together, these results demonstrate that higher PA levels are associated with prolonged CD63 decay during MVE fusion and that experimental PA elevation increases the prevalence of these events.

## DISCUSSION

### PLD1 is on MVEs and necessary for exosome release in A549 cells

In this work, we demonstrate that PLD1 is needed for exosome secretion in A549 cells where the inhibition of PLD1 dramatically reduced the number of secreted exosomes (Figure 1C). It is worth noting that we used the PLD1-selective inhibitor VU0155069 (IC_50_ values are 46 and 933 nM for PLD1 and PLD2 respectively) (69); and the pan-PLD inhibitor FIPI (reported cellular IC50 ≈ 1–25 nM toward PLD1/2)(70). At 500 nM VU0155069, PLD2 inhibition is unlikely, whereas FIPI inhibits both isoforms; effects of inhibition with VU0155069 therefore supports a PLD1-specific requirement for exosome release. To further verify the requirement of PLD1 for exosome secretion, PLD1 knockdown experiments were performed and show a similar reduction (Fig 1D). These observations demonstrate that PLD1 activity is required for exosome secretion, and the requirement for PLD1 activity is conserved across several model cancer cell lines (3T1, MDA-MT, ova, RED) (20, 21).

Using immunofluorescence, we observed endogenous PLD1 on CD63^+^ vesicles (Figure 1A–B). Colocalization of PLD1 with CD63 has been reported in multiple cell lines (23–25, 27, 63). Consistent with this localization, PLD1 may contribute to MVE exocytosis through local PA production. In stimulated secretion, PLD1 generates PA at exocytic sites (26, 28, 29, 35, 36, 71, 72). Mechanistically, PA can (i) bind the positively charged region of syntaxins near the membrane to modulate SNARE function ; (ii) activate type I PI4P 5-kinases to elevate local PI(4,5)P_2_ and recruit exocytic effectors (73); and (iii) stabilize negative curvature (32, 36, 38, 39, 74). Each mechanism would favor MVE–plasma membrane fusion when PLD1 is active at the vesicle–plasma membrane interface. PLD1 localization to MVEs therefore positions it to contribute to regulation of exosome secretion paralleling its role in stimulated exocytosis (26, 51, 62, 75)

To map when and where PLD1 is required in the exosome secretion process, we considered the sequential steps of exosome secretion as: (1) MVE trafficking, (2) MVE docking, and (3) MVE fusion. To visualize pre-fusion stages, we used a CD63 construct with pHmScarlet—rather than pHluorin—to improve visualization of MVEs prior to fusion (Figure S2; (50)); pHmScarlet is not completely quenched in the acidic MVE lumen, allowing dim but detectable signal before fusion. In our experience, MVEs containing CD63-pHmScarlet appear at ∼10% of their maximum intensity. We then focused on PLD1 and PA at each step to build a working model.

1. *MVE trafficking:* we assessed trafficking in two ways. (i) By real-time imaging of single CD63-pHmScarlet–labeled vesicles near the plasma membrane together with GFP-PLD1 or GFP-PASS (PA sensor) (Figures 2A and 4A). PLD1 is present on moving vesicles near the cell surface (Figure 2A), whereas PA is detectable but increases after docking (Figure 4A–B, E). Thus, moving vesicles carry PLD1 and some PA, with PA rising upon docking. (ii) By counting the number of CD63+ vesicles present at the plasma membrane to indicate whether they are trafficked to the cell surface (Figure 3B). PLD1 inhibition caused a small but significant reduction in membrane proximal MVEs (Figure 3B). These data indicate that PLD1 activity is not required for the bulk of transport to the cell surface but plays a small role, possibly like that observed in dense core vesicles where PLD1 inhibition slightly slowed the rate of vesicle motion (26, 28, 28, 72, 76).
2. *MVE docking:* To visualize the presence of PLD1 and PA during docking, we used CD63pHmScarlet alongside with GFP-PLD1 or GFP-PASS. Prior to docking, PLD1 and PASS are enriched at the docking location (Figures 2B, 2E, 4B, 4E), relative to cytoplasmic GFP controls (Figure S3B, 2E, 4D). Once the vesicle docks, both PLD1 and PASS increase slowly over time, however cytoplasmic GFP does not (Fig 2E, 4E). Based on neuroendocrine exocytosis, PLD1 and PA likely support vesicle docking. One plausible mechanism involves the SNARE machinery: PA binds syntaxin-1 (Syx1)(72, 77, 78), PLD1 colocalizes with Syx1 (26), and Syx1 participates directly in dense-core vesicle docking(40, 41, 79, 80). In our past work, PA accumulated over tens of seconds at docked vesicles in PC12 cells (26), and similarly we observe PA accumulation at docked MVEs. Whether this PA resides primarily on the plasma membrane, the vesicle membrane, or both remains unresolved. The intensity of GFP-PASS increases during docking but is lower immediately before fusion. We interpret this decline in PASS as probe displacement. It is possible that PA-binding proteins, such as syntaxins, compete with PASS for PA-rich sites (72, 77, 81); Syx2/3/4 have been implicated in exosome secretion (17, 44, 82). Alternatively, PA may decline just prior to fusion as other factors stabilize the docked state. Despite PLD1 inhibition, MVE surface delivery is largely preserved (Fig. 3B) yet the probability that a docked vesicle fuses is diminished (Fig. 3A). If we consider the 55% reduction in fusion, we expected to see a rise in docked vesicles if fusion was the only step blocked. Instead, a 20% reduction in membrane proximal CD63+ vesicles was observed, suggesting an alternate pathway. CD63+ vesicles arrive at the plasma membrane but then do not secrete. Some of these vesicles likely become lysosomes (Fig 3C) whereas other pathways could include undocking or other alternatives. Thus, PLD1 catalytic activity is likely required to stabilize docking and confer fusion competence; when curtailed, MVEs do not fuse and are biased toward lysosomal routing.
3. *MVE fusion:* The fusion of single MVE was measured by observing the unquenching of CD63-pHmScarlet followed by a radial spread of the fluorescence (Fig 2C, 4C). Concurrently, PLD1 or PASS was measured with GFP-tagged probes. As we and others have previously reported, CD63 fusion events do not contain lysosome markers (46, 83–85). These fusion events are also not membrane trafficking vesicles as they are unaffected by Brefeldin A treatment (46). Therefore, most, if not all, fusion events we identified in this work were MVE fusion events.

Given that PLD inhibition (PLD1-selective and pan-PLD; (Figure. 1) as well as PLD1 knockdown reduce secreted exosomes (Figure. 3A), it is likely that PLD1 produces PA during MVE exocytosis. The main enzymatic activity of PLD1 is conversion of PC to PA (86, 87), which is hypothesized to stabilize negative curvature(28, 29, 31, 33, 88, 89). Therefore, we hypothesized that PA may stabilize membrane curvature at the exocytic site, as reported for dense core vesicle secretion (28, 29, 35, 88), and/or support ILV formation, another curvature-rich process (41, 53, 83). Others have reported that lipids favoring negative curvature tend to stabilize late exocytic intermediates or limit rapid expansion at the release *site* To explore this possibility, we examined how the post-fusion fluorescence loss rate related to PA content (Fig. 5). After clustering fusion decays by release half-time (t_1_/_2_; Fig. 5A–B), we measured the PASS intensity in each cluster (Fig. 5C). The rate of release was inversely correlated with PASS, with higher PASS associated with slower CD63 decay. This supports the hypothesis that PA stabilizes curvature, or recruit proteins that do so, and this restricts fusion pore expansion and the release of content.

Although the GFP-PASS probe is used as a PA marker, it is worth noting that the binding of the probe depends on the accessibility of the PA headgroup and is influenced by other lipids (28, 35, 43, 90). As an 18-amino-acid, amphipathic helix with a positively charged face, the probe’s specificity is primarily due to electrostatic interactions and an ability to embed in membranes (28, 32, 74, 81, 90). In vitro studies show that a dimeric PASS binds PA more strongly in phosphatidylethanolamine (PE)-rich membranes— conditions that stabilize PA in the −2 charge state (vs −1)—and on curved membranes. PASS can also bind PI(4,5)P_2_ and other negatively charged lipids (43, 91, 92). However, no other labeling options compatible with high-speed (100 ms), time-resolved, live-cell microscopy exist for PA. Therefore, interpretations of PASS data are likely influenced by the presence of other lipids and the local lipid nano-environment. Further studies can elucidate the roles of other lipids and membrane shape.

### Lysosomal routing as an alternate to fusion

When fusion is blocked, the number of membrane-proximal, catalytically active lysosomes increases (Fig. 3C). Consistent with this, when membrane conditions do not support fusion, MVEs appear to be diverted to lysosomes and PLD1 influences late endosome/MVE fate(26, 93). Upon inhibition, a small fraction of CD63^+^ vesicles at the proximal membrane decreases by ∼20% (Fig. 3B), and a similar ∼23% increase in lysosomes is observed (Fig. 3C). Although this shift is modest, PLD1 activity is necessary for fusion (Fig. 1C–D), and disruption biases vesicles toward the lysosome pathway. It is possible that a longer incubation period could lead to more substantial changes. In Fig. 1C, cells were treated with inhibitors throughout the sEV collection period (2 days), whereas for fusion events (Fig. 3A), CD63^+^ vesicle density (Fig. 3B), and lysosome density (Fig. 3C), cells were acutely treated (30 min).

## Conclusion

PLD1 activity is necessary for exosome secretion. However, due to the strong reduction in fusion upon knockdown or inhibition of PLD1, it was challenging to gather enough events to determine exactly when and where PLD1 acts. Nevertheless, our data support: (1) endogenous PLD1 on CD63^+^ MVEs (fixed-cell colocalization; Fig. 1A–B); (2) PLD1 is required for fusion (knockdown reduces single-vesicle fusion; Fig. 1D); (3) PLD1 activity is required (inhibitors reduce fusion; Fig. 1C); (4) PLD1 inhibition increases membrane-proximal lysosomes (Fig. 3D); (5) PA accumulates as MVEs remain docked (PASS increases over time; Fig. 4B), though whether this reflects PLD1 activity at the docked MVE and/or PA binding at the docking site remains unresolved. (Figure 4B); and (6) higher PASS/PA correlates with slower CD63 decay (Fig. 5), consistent with PA stabilizing fusion pore.

## Supporting information

Supplemental Figures_Nguyen

## Abbreviations

PLD1: Phospholipase D1
PA: Phosphatidic Acid
PABD: PA Binding Domain
MVE: Multivesicular Endosome.

## AUTHOR CONTRIBUTIONS

MTN performed experiments and aided with experimental design, analysis, and writing of the final draft. BLB performed experiments and aided with experimental design, analysis, and initial drafts. ACA performed confocal microscopy, MLH performed TIRF experiments and analyzed data, CJ cloned the CD63-pHmScarlet probe, MKK designed experiments, obtained funding for the work, analyzed data, and drafted the paper.

## ACKNOWLEDGEMENTS

This work was funded by the National Science Foundation Chemistry of Life Processes (grant # 2325227)

