## Supplemental Figures_Nguyen for "Phospholipase D1 and Phosphatidic Acid are required for MVE Fusion and Exosome Secretion"

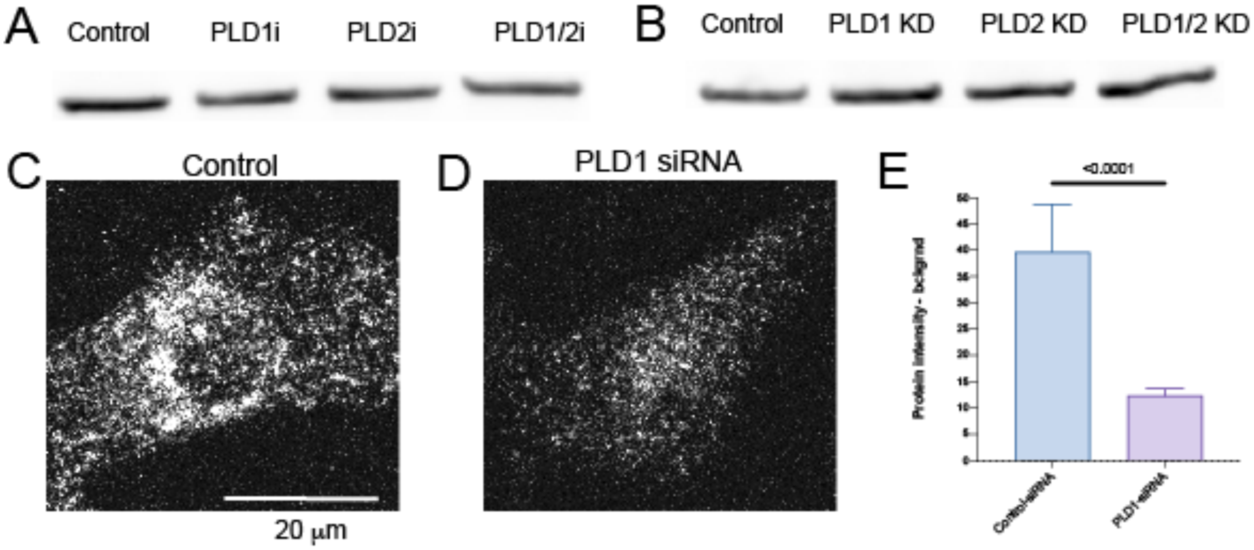


**Supplemental Figure 1: Control experiments for sEV collection and NTA measurements** (A, B) Representative actin (42 kDa) blots shown as loading controls for the indicated pharmacological inhibition and siRNA-mediated knockdown conditions. The WB band of actin was used to account for any reduction in cells that may have occurred when treated with PLD inhibitors. (C, D) Immunofluorescent confocal microscopy images of A549 cells transfected with control siRNA (C) or PLD1 siRNA (D). Scale bar: 20.0 µm. (E) Quantification of PLD1 intensity from confocal images shows a significant reduction in PLD1-siRNA–treated cells ($p < 0.0001$**),** likely underestimated due to ~35% transfection efficiency. Data are mean ± SEM from three independent experiments.

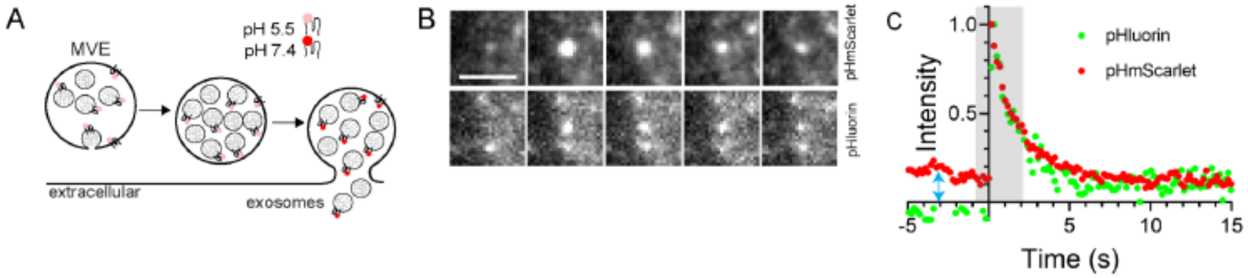


**Supplemental Figure 2: CD63-pHmScarlet is a probe for measuring MVE fusion and can be observed prior to fusion, more so than CD63-pHluorin.** (A) Schematic illustrating how intraluminal vesicles (ILVs) labeled with pH-sensitive CD63 probes are quenched in the acidic lumen of multivesicular bodies (MVBs) and become unquenched upon fusion with the plasma membrane, exposing the probe to the neutral extracellular environment. (B) Representative montage images showing a single fusion event for CD63-pHmScarlet (top) and CD63-pHluorin (bottom), spanning 10 frames before to 20 frames after fusion onset (the shaded window in panel C). (C) Averaged CD63 intensity traces for CD63-pHmScarlet (red) and CD63-pHluorin (green) aligned to fusion onset. CD63-pHmScarlet exhibits a higher pre-fusion baseline due to incomplete quenching in acidic compartments, enabling visualization of docking vesicles before fusion, an advantage for tracking docking behavior in live-cell imaging.

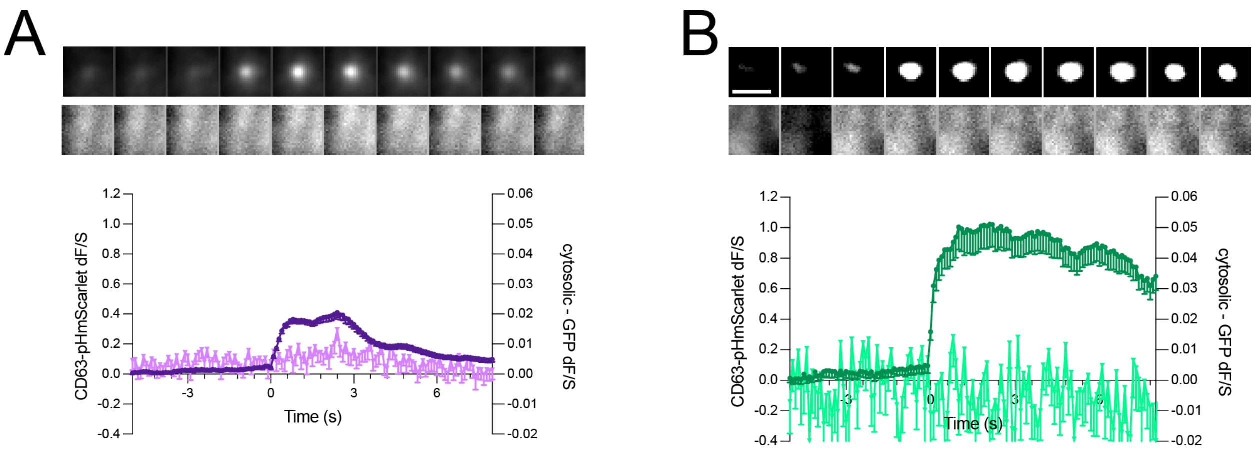


**Supplemental Figure 3: CD63-pHmScarlet and cytosolic-GFP visiting and docking average traces.** (A–B) Representative traces of CD63 (darker shade) and cytosolic GFP (lighter shade) are shown for visiting (A, purple), docking (B, green). CD63 (top row) and GFP (bottom row) image montages span from 49 frames before onset to 80 frames after. Each montage frame represents a 15-frame average (1.5 seconds). Examples were selected from a dataset of 161 visiting, 33 docking. (Scale bar: 2.0μm).

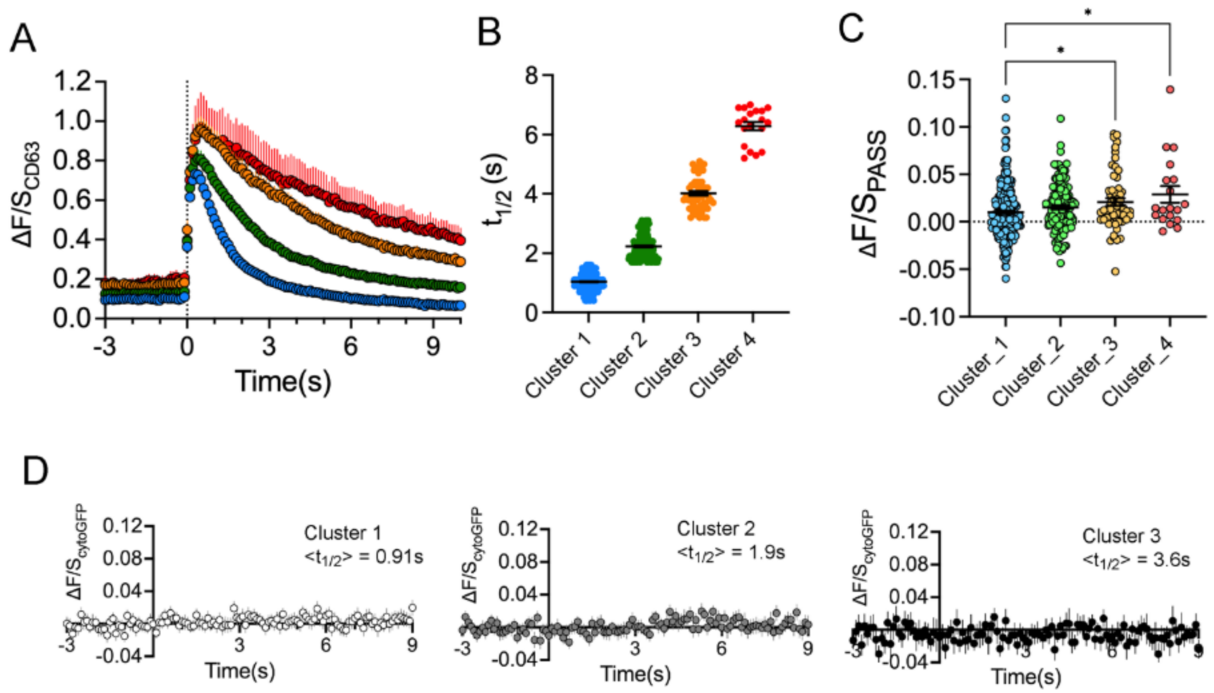


**Supplemental Figure 4: Cluster-resolved CD63 decay kinetics.** (A) Per-cluster mean CD63-pHmScarlet intensity traces aligned to fusion onset (t = 0) for four clusters: Cluster 1 (blue, n = 252), Cluster 2 (green, n = 141), Cluster 3 (orange, n = 57), and Cluster 4 (red, n = 19). Traces are averaged across events within each cluster; shading denotes ±SEM where shown. (B) Decay duration (s) for each cluster in (A), summarizing the distribution of fusion-decay times. "Decay duration" is defined as the time from the event-specific CD63 peak to the half-maximum level ($t_{1/2}$). (C) Mean single-event ΔF/S trajectories of PASS (PA sensor) computed from the CD63 peak to that event's $t_{1/2}$, grouped by cluster. (D) Mean cytosolic-GFP intensity traces (negative control), processed and aligned identically, shown for three clusters, demonstrating no event-locked modulation.

**Supplemental Table 1: Data Table for Figure 3 - Fusion events per minute for inhibited treatments.** Inhibitors were purchased twice and results were reproducible from one batch to the next as well as from one year to the next. The rate of fusion from control cells varies substantially over time for reasons not well understood. Therefore, all experiments are paired with data take on the same day. The fraction of control is plotted in Figure 3A. Events per minute is the raw data.

| | **Events per minute** | | | **Fraction of control** | |
|---|---|---|---|---|---|
| | **Control** | **$PLD1_i$** | **$PLD1/2_i$** | **$PLD1_i$** | **$PLD1/2_i$** |
| **Exp#1 2024** | 1.26 | 0.57 | 0.80 | 0.45 | 0.63 |
| **Exp#2 2024** | 1.60 | 0.73 | 1.17 | 0.46 | 0.73 |
| **Exp#3 2024** | 1.47 | 0.96 | 0.68 | 0.65 | 0.46 |
| **Exp#1 2025** | 2.57 | 0.21 | 1.39 | 0.08 | 0.54 |
| **Exp#2 2025** | 2.95 | 1.65 | 1.39 | 0.56 | 0.47 |
| **Exp#3 2025** | 2.94 | 1.51 | 1.10 | 0.51 | 0.37 |
| **Exp#4 2025** | 2.67 | 1.12 | 0.28 | 0.42 | 0.11 |

**Supplemental Table 2: Data Table for Figure 3 - Fusion events per minute for siRNA treatments.** The Fraction of control is plotted in Figure 3A. Events per minute is the raw data.

| | **Events per minute** | | | **Fraction of control** | |
|---|---|---|---|---|---|
| | **Control** | **$PLD1_{KD}$** | **$PLD1/2_{KD}$** | **$PLD1_{KD}$** | **$PLD1/2_{KD}$** |
| **Exp #1** | 1.64 | 0.24 | 0.25 | 0.15 | 0.15 |
| **Exp #2** | 1.00 | 0.63 | 0.23 | 0.63 | 0.23 |
| **Exp #3** | 1.63 | 0.60 | 0.30 | 0.37 | 0.18 |
| **Exp #4** | 1.93 | 0.80 | 0.83 | 0.41 | 0.43 |
